# Defining the molecular impacts of Humalite application on field-grown wheat (*Triticum aestivum* L.) using quantitative proteomics

**DOI:** 10.1101/2024.12.18.629253

**Authors:** Lauren E. Grubb, Mohana Talasila, Linda Y. Gorim, R. Glen Uhrig

**Affiliations:** Department of Biological Sciences, University of Alberta, Edmonton, AB, CAN; Department of Agriculture, Food and Nutritional Science, University of Alberta, Edmonton, AB, CAN; Department of Biochemistry, University of Alberta, Edmonton, AB, CAN

**Keywords:** Biostimulants, BoxCarDIA, humic acids, Humalite, quantitative proteomics

## Abstract

Increasing global food production demands have resulted in increased fertilizer usage, causing detrimental environmental impacts. Biostimulants, such as humic substances, are currently being applied as a strategy to increase plant nutrient-use efficiency and minimize environmental impacts within cropping systems. Humalite is a unique, naturally occurring coal-like substance found in deposits across southern Alberta. These deposits contain exceptionally high ratios of humic acids (>70%) and micronutrients due to their unique freshwater depositional environment. Humalite has begun to be applied to fields based on scientific data suggesting positive impacts on crop growth, yield and nutrient usage; however, little is known about the underlying molecular mechanisms of Humalite. Here, we report a quantitative proteomics approach to identify systems-level molecular changes induced by the addition of different Humalite application rates in field-grown wheat (*Triticum aestivum* L.) under three urea fertilizer application rates. In particular, we see wide-ranging abundance changes in proteins associated with several metabolic pathways and growth-related biological processes that suggest how Humalite modulates the plant molecular landscape. Overall, our results provide new, functional information that will help better inform agricultural producers on optimal biostimulant and fertilizer usage.

## 1. INTRODUCTION

Wheat (*Triticum aestivum* L.) represents a critically important cereal crop worldwide, acting as a staple food for over 2.5 billion people^1^. To increase grain yield, large quantities of nitrogen (N), phosphorus (P), and potassium (K) fertilizers are applied to fields. Nitrogen is an important nutrient and is a major component of important plant compounds such as amino acids, proteins, nucleic acids and chlorophyll^2^. Although important for driving agricultural yield, excessive N fertilizer application has detrimental impacts on the environment. On average, only 40-70% of applied N fertilizer is biologically available for crop plants^3^, with the excess contributing to agricultural runoff, soil acidification, nitrate leaching into groundwater, and greenhouse gases^4–7^. Therefore, sustainable solutions for increasing crop yield while minimizing environmental harm are warranted.

A recent strategy to increase nitrogen use efficiency in field crops is the use of biostimulants, such as humic substances, which are the product of decomposition of plant and animal materials^8^. Humic substances are composed of approximately 80% soil organic matter, and are made up of humic acids, fulvic acids and humins, depending on their solubility in water, acidic or alkaline solutions^9,10^. They are reported to improve the yield of crop plants, including maize (*Zea mays)*^11–16^, wheat (*T. aestivum)*^17–19^, rice (*Oryza sativa)*^20^, foxtail millet (*Setaria italica)*^21,22^, soybean (*Glycine max)*^23^, chickpea (*Cicer arietinum)*^24^, cucumber (*Cucumis sativus)*^25^, barley (*Hordeum vulgare)*^26^, canola (*Brassica napus)*^27,28^ and pea (*Pisum sativum)*^29^. Beneficial impacts on plant physiology with humic acid application have been reported, including modification of root system architecture^13,27,30^ and improved nutrient use efficiency^27,31^. Furthermore, humic substances have been reported to increase plant performance under abiotic stress such as drought^17,21,22,32^, salt^12,33,34^, heat^35^ and heavy metals^36,37^.

Several studies have also found that humic acid in combination with synthetic fertilizer, such as urea, can further increase crop yields^38–40^. Humic acid urea, an organic-inorganic compound fertilizer consisting of a mixture of conventional urea with humic acids, has shown promise for increasing crop yield, biomass, and nitrogen uptake^41–43^. However, little is known about the plant molecular interplay of humic acids in combination with urea.

Despite a variety of studies examining the phenotypic effects of humic acid application on crops, there remains a significant lack of information on how it affects the plant molecular environment. Several transcriptomic and microarray analyses^27,31,44,45^ have identified changes in molecular processes such as plant C/N/S metabolism, photosynthesis and respiration, cell cycle, cytoskeleton, auxin signaling, response to stress and ion / water transporters as potential drivers in the beneficial observations with humic acid treatment. A few proteomic studies have been performed; however, humic acid studies to date have all been conducted under controlled environment conditions and involved hydroponic systems to examine roots^13,46,47^ or cell culture^48^, using a variety of humic acids from different sources. Additionally, these molecular studies were typically performed at earlier developmental timepoints than our study and examined immediate responses with humic acid addition to hydroponic media, or with foliar application. Reports of the transcriptome at multiple timepoints also found more fast-responsive genes than those at later timepoints^27,44^, however transcriptional responses are known to be a poor indicator of protein-level changes that may have functional outcomes^49,50^. Given these experimental approaches, there is a pressing need to understand the molecular responses of crops to humic acid application under field conditions.

Humalite is a humic substance that is a naturally occurring biostimulant, found in large deposits in Southern Alberta, which contains a uniquely high concentration of humic acids (>70%) and very low amounts of nutrients and heavy metals due to its unique freshwater depositional environment. Farmers have begun to apply Humalite to their fields in Western Canada, with little knowledge on appropriate rates and impacts on crops. However, recent studies conducted using Humalite^18,26,29,51^ under controlled greenhouse conditions, report enhanced growth, grain yield, protein content and nitrogen use efficiency^18^. Correspondingly, as part of a larger field study^19^, we investigated the molecular impacts in wheat of Humalite application under no, half, and full recommended urea application rates using quantitative proteomics. We report striking changes in the abundance of proteins associated with plant C/N metabolism, carbohydrate metabolism, protein folding, heat responses, and vesicle transport. We additionally report differences in the regulation of these processes with the addition of Humalite dependent on the urea application level. Our results contribute increased precision in our molecular understanding of the biological processes underpinning the physiological impacts of Humalite addition in combination with urea fertilizer, offering to more robustly inform best practices for use of Humalite in crop fields.

## 2. MATERIALS AND METHODS

### 2.1 Plant growth and tissue harvesting

Wheat (*Triticum aestivum* L.) field trials were established at St. Albert Research Station, University of Alberta (53.6929508, −113.6353861) in a split-plot design as detailed for the year 2023 of a 3-year study^19^, with soil described as Luvic Chernozem (IUSS Working Group WRB, 2022) and soil texture as silty clay loam (sand 6%, silt 56%, and clay 38%). The main plot had urea fertilizer applied at three rates based on soil tests: no urea, half the recommended amount of urea (112 kg ha^-1^), and full recommended urea application rate (244 kg ha^-1^). The subplots had Humalite (supplied by WestMet Ag, Hanna, Alberta, Canada), specifically from the holdings of the Sheerness Coal Mine (recently: WestMet Ag) applied at four rates plus a zero Humalite control: 0, 56, 112, 224, 448 kg ha^-1^. Humalite and urea were side-banded at seeding. At BBCH stage 31, 3-4 wheat leaves were rapidly collected from multiple random plants within each plot, 3 biological replications per plot. Each treatment had 3 plots within the field set-up for a total of n = 9 total replicates for each treatment. Harvested leaves were flash-frozen in liquid nitrogen and transported back to the laboratory on dry ice followed by storage at −80 °C until processing.

### 2.2 Tissue processing, protein extraction and trypsin digestion

Under liquid N_2_, wheat leaves were ground to a fine powder using a Genogrinder (SPEX SamplePrep) at 1,200 rpm for 30 seconds repeated until fully ground. Ground material was aliquoted into 100 mg fractions followed by extraction in a solution of 1:2 (w/v) 50 mM Tris-HCl pH 8.0, and 4% (w/v) SDS. Samples were briefly vortexed and then placed in a table-top shaking incubator (Eppendorf ThermoMixer F2.0) set to 95 °C, shaking at 1,000 rpm for 10 min followed by 5 min shaking (same speed) at room temperature. Samples were then centrifuged at 20,000 x g for 5 min to clarify extracts, and the supernatant was transferred to a fresh 1.5 mL Eppendorf tube. Protein concentration was measured by Bradford assay and normalized to 23 mg in 250 µL extraction buffer. Samples were subsequently reduced with 10 mM dithiothreitol (DTT) at 95 °C for 5 min, cooled to room temperature, and then alkylated with 30 mM iodoacetamide (IA) in the dark for 30 min at room temperature without shaking. To stop the reaction, 10 mM DTT was added to each sample, followed by a brief vortex and then incubation for 10 min at room temperature without shaking. Samples were digested using sequencing grade trypsin (V5113; Promega) at 37 °C overnight as previously described^52^ without deviation. The generated peptide pools were acidified with formic acid (FA) to a final concentration of 0.5% (v/v) prior to desalting with ZipTip C18 pipette tips (ZTC18S960; Millipore). Samples were then dried and resuspended in 3% (v/v) ACN / 0.1% (v/v) FA prior to MS analysis.

### 2.3 LC-MS/MS analysis

A total of 2 µg resuspended peptide was eluted into the mass spectrometer using an Easy-nLC 1200 system (Thermo Fisher Scientific) and an ES906 (Thermo Fisher Scientific) Easy-Spray PepMap C18 analytical column as previously described^53^, using −30, −50, −70CVs. Peptides were analyzed using FAIMSpro enabled Fusion LUMOS Orbitrap mass spectrometer (Thermo Fisher Scientific) in BoxCarDIA mode previously described^53^ without deviation.

### 2.4 Data analysis

Acquired BoxCarDIA data was analyzed using a library-free DIA approach in Spectronaut v18 (Biognosys AG) under default settings. Data were searched against the Ensembl Plant *Triticum aestivum* database (https://plants.ensembl.org/Triticum_aestivum). Search parameters included the following: trypsin digest allowing two missed cleavages, fixed modifications (carbamidomethyl (C)), variable modifications (oxidation (M)) and a peptide spectrum match, peptide and protein false discovery threshold of 0.01. Significantly changing differentially abundant proteins were determined and corrected for multiple comparisons (Bonferroni-corrected p-value <0.05; q-value).

### 2.5 Bioinformatics

Gene Ontology (GO) analyses were performed with the Database for Annotation, Visualization and Integrated Discovery (DAVID; v 6.8; https://david.ncifcrf.gov/home.jsp), using significantly changing proteins for the indicated categories as a search list and the *Triticum aestivum* proteome as background. Significance threshold was set to a p-value <0.01. GO dot plots and volcano plots were visualized using *R* version 4.3.1 and the *ggplot2* package. For metabolic pathway analysis and association network analysis, wheat gene identifiers were converted to Arabidopsis gene identifiers (AGI) using UniProt (https://www.uniprot.org/). Metabolic pathway analysis enrichment was done using the Plant Metabolic Network (PMN; https://plantcyc.org/) with significance determined by a Fisher’s exact test threshold p-value of <0.01. Association network analysis was performed using the STRING-DB app (https://string-db.org/) in Cytoscape (version 3.10.1; https://cytoscape.org/) with a minimum edge threshold of 0.7 and the CIRCOS enhancedGraphics plugin (version 1.5.5; https://apps.cytoscape.org/apps/). Completed figures were assembled using Affinity Designer (v. 2.4.1; https://affinity.serif.com/en-us/designer/).

## 3. RESULTS

### 3.1 Effect of Humalite on the wheat proteome

To investigate the molecular effects of Humalite and its interplay with urea fertilizer, we employed a split-plot growth design in the field at the University of Alberta Research Station in St. Albert (Alberta, Canada), as part of a larger study field study^19^, as described in the Materials and Methods section (**Figure 1a**). At BBCH stage 31, multiple leaves from randomly selected plants were sampled for a total of three replicates per plot, each in three different plot areas for a total n = 9 total replicates for each treatment. We next used high-resolution LC-MS/MS to quantify changes in the wheat proteome across different Humalite and urea fertilizer application rates. In total, we quantified 6,035 protein groups across all experimental treatments (**Figure 1b**; **Supplemental Table 1).** Next, we queried our data for proteins showing a significant change in abundance upon addition of Humalite in comparison to the ‘No Humalite’ treatment at each of the three urea levels (Log2FC < −0.58 or > 0.58, q-value < 0.05; **Figure 1b**). Interestingly, with both half and full urea amounts, the number of significantly changing proteins seemed to increase with increasing Humalite, with the most significantly changing in abundance under the ‘Half Urea’ treatment with 448 kg ha^-1^ Humalite. There was also considerable overlap in the significantly changing proteins within the urea treatments with any level of Humalite application. Here, we found a total of 478 proteins significantly changing in abundance with Humalite between all three urea treatments (**Figure 1c**). We next examined the general trends in the observed protein abundance changes; whether up- or down-regulated with Humalite under the urea application regimes. Strikingly, volcano plots indicated that the addition of Humalite resulted in significantly more up-regulated proteins under the ‘No Urea’ treatment, but more significantly down-regulated proteins under the ‘Half Urea’ or ‘Full Urea’ application rate (**Supplemental Figure 1**).

**Figure 1.**
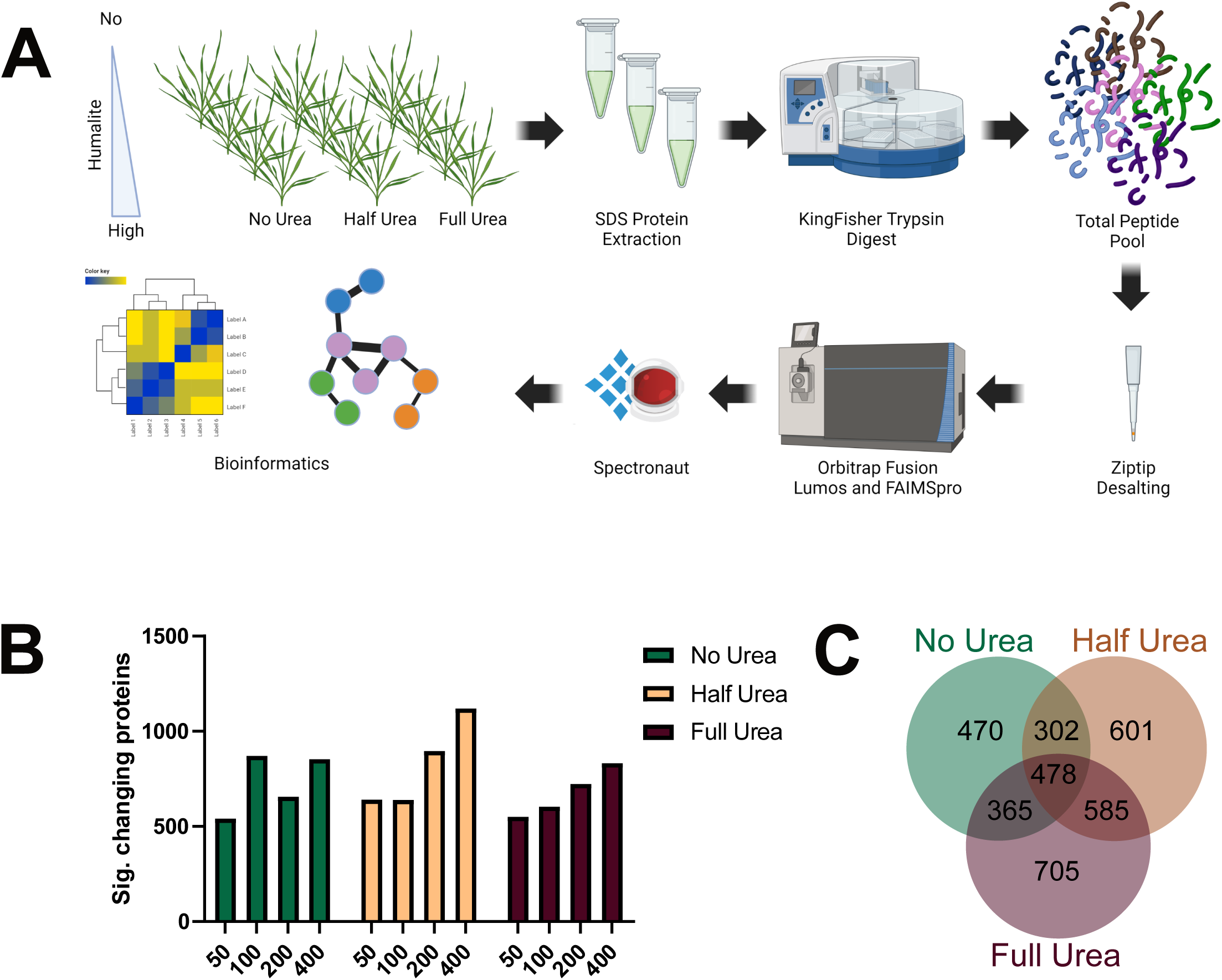
Experimental design and summary of proteome data A) Experimental workflow schematic. Wheat seeds were sown and grown at the University of Alberta Research Station, St. Albert, Alberta on soil with no, half and full recommended value with 0, 50, 100, 200, or 400 kg*ha-1. Several leaves from randomized plants were harvested at the point at which 80% of the field had germinated (n=3 from each of 3 plot positions for a total of 9 reps). Proteins were extracted, digested and desalted and total proteome was determined using LC-MS/MS with FAIMSpro and BoxCarDIA acquisition. B) Number of significantly changing proteins (q-value <0.05; Fold change >1.5) with no urea (green), half urea (beige) and full urea (burgundy) with addition of the indicated Humalite (50, 100, 200, 400 kg/ha). C) Venn diagrams of overlap of significantly changing proteins overlap under fertilizer regimes with any amount of Humalite addition.

### 3.2 Effects of Humalite on plant metabolism

Given the observed trends in up- and down-regulated proteins, we proceeded to investigate the biological processes that are up- or down-regulated to identify trends in the addition of Humalite across the three urea levels. To do this, we generated a heatmap using correlation clustering of the Log2FC values for proteins significantly changing across at least two different Humalite levels within a single urea application rate (**Figure 2a**). This led to the identification of three clusters of similarly significantly changing proteins that were dependent on urea application rate (**Figure 2b**). We next performed Gene Ontology (GO) analysis on each cluster to determine trends in the biological processes associated with Humalite addition across different urea application rates (**Figure 2c**). Cluster 1 contains the greatest number of significantly changing proteins (976 proteins) up-regulated under ‘No Urea’ and significantly down-regulated under ‘Half Urea’ and / or ‘Full Urea’. This cluster was enriched for biological processes such as: Protein Transport, Iron-Sulfur Cluster Assembly, Pentose-Phosphate Shunt, Fatty Acid Beta-Oxidation, Starch Biosynthetic Process, Lipid Metabolic Process, Carbohydrate Metabolic Process and Response to Reactive Oxygen Species. Cluster 2 (357 proteins) corresponded to proteins exhibiting no change under the ‘No Urea’ treatment, but a decrease under ‘Half Urea’ and increase under ‘Full Urea’ treatments. This cluster was enriched for Chromosome Condensation, Glucose 6-phosphate Metabolic Process, Intracellular Glucose Homeostasis, Lignin Biosynthetic Process, Response to Oxidative Stress and Proteolysis. Finally, Cluster 3 (530 proteins) consists of proteins significantly up-regulated under ‘Half Urea’, but down-regulated under the ‘No Urea’ and ‘Full Urea’ treatments and was significantly enriched for proteins involved in Protein Folding and Electron Transport Chain.

**Figure 2.**
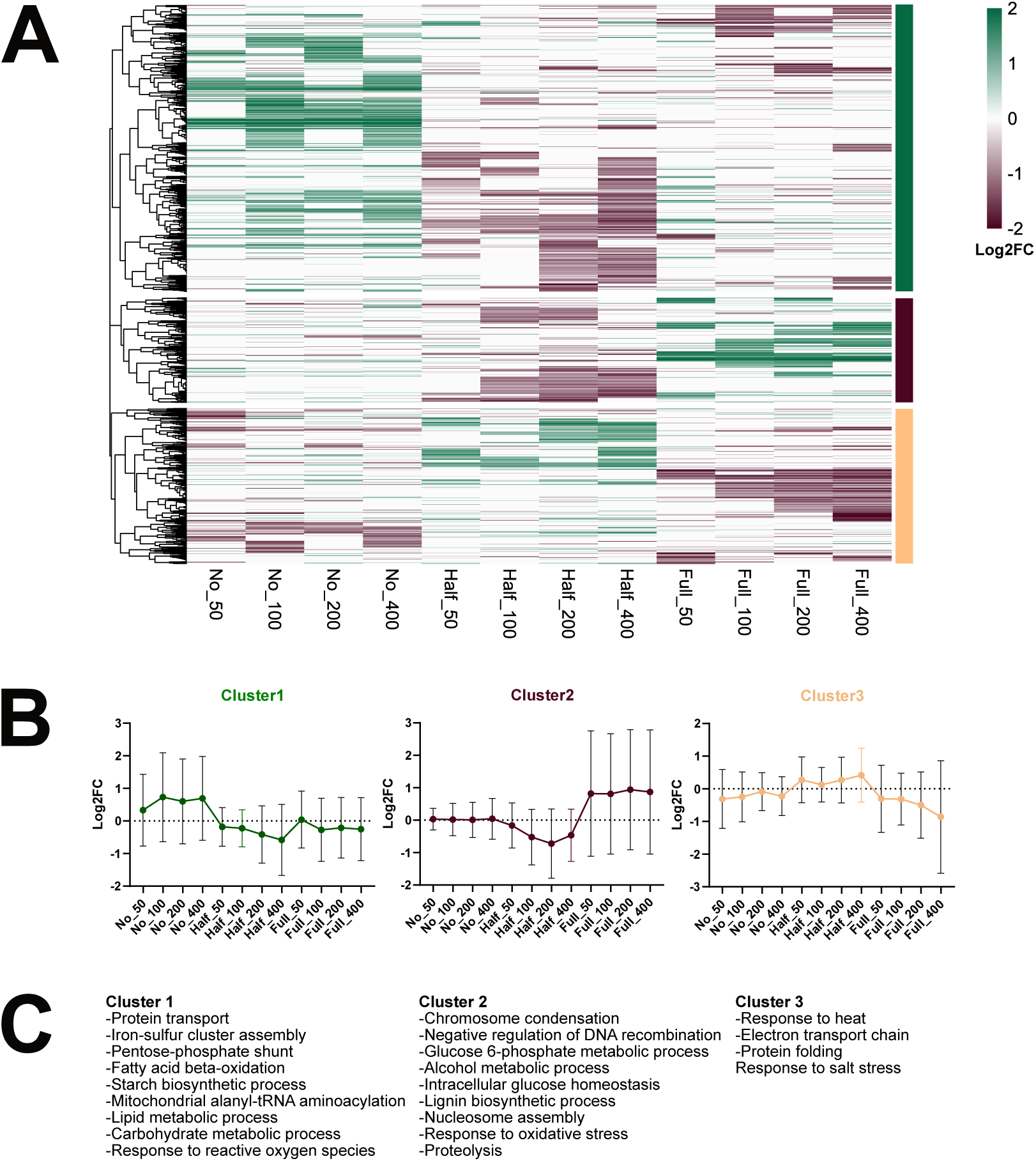
Proteome-level changes with Humalite addition under urea application regimes A) Heatmap of significantly changing proteins with Humalite application under the urea application regimes (Log2FC scale in comparison to no Humalite). Correlation clustering was performed and identified 3 main clusters. B) Breakdown of identified clusters over the Humalite and urea application regimes. y-axis shows Log2FC in comparison to no Humalite under the indicated urea application rate. C) Selected enriched GO biological processes of each cluster (p<0.01).

Given that our GO results suggest that Humalite has major impacts on plant metabolism, we aimed to further elucidate plant metabolic responses to Humalite application, through the Plant Metabolic Network (PMN; https://plantcyc.org/; **Supplemental Table 2**). Here, we undertook a metabolic pathway enrichment analysis using the significantly changing proteins (Log2FC < −0.58 or > 0.58, q-value < 0.05). As the PMN lacks *Triticum aestivum* resources, we first identified Arabidopsis orthologs for the significantly changing proteins using UniProt (https://www.uniprot.org/). We were able to convert 70.6% (4259/6035) of the wheat gene identifiers for the significantly changing proteins to Arabidopsis gene identifiers (AGI) for our metabolic pathway analysis. Consistent with our GO analysis, for Cluster 1 we observe changes in metabolic pathways associated with Aerobic Respiration, Starch Biosynthesis, and Fatty Acid Degradation, and additionally identify Glutamate-Glutamine Shuttle associated with N metabolism. For Cluster 2, we find metabolic pathways associated mostly with Carbohydrate Metabolism, while Cluster 3 was enriched in metabolic pathways for N Assimilation, Amino Acid Biosynthesis, and Fatty Acid Beta-Oxidation.

### 3.3 Network analysis identifies striking differences in wheat proteome upon Humalite application with varying levels of urea fertilization

To further resolve our GO and metabolic pathway analyses, we next performed an association network analysis using STRING-DB (https://string-db.org/). To best contextualize our results, this analysis utilized proteins significantly changing in abundance at two or more Humalite application rates within each level of urea application and using a STRING-DB edge threshold > 0.7 (**Figure 3**). Consistent with our previous analyses, we observe striking differences in the regulation of processes dependent on the level of urea fertilizer, with the ‘No Urea’ treatment possessing mostly up-regulated proteins, and the ‘Half Urea’ and ‘Full Urea’ treatments mostly down-regulated proteins. This analysis revealed clear clusters of proteins consistently measured across different treatments that are associated with carbohydrate metabolism, fatty acid metabolism, nitrogen metabolism, lignin metabolism and oxidative stress response that were generally up-regulated in the ‘No Urea’ treatment and down-regulated with the ‘Half Urea’ and ‘Full Urea’ treatments. We additionally resolved clusters of up-regulated proteins associated with growth and development under the ‘No Urea’ treatment, as well as a notable number of significantly changing proteins associated with photosynthesis and chlorophyll metabolism across all urea treatments. Overall, these results indicate the addition of Humalite induces substantial changes in biological processes associated with metabolism and plant response to stress.

**Figure 3.**
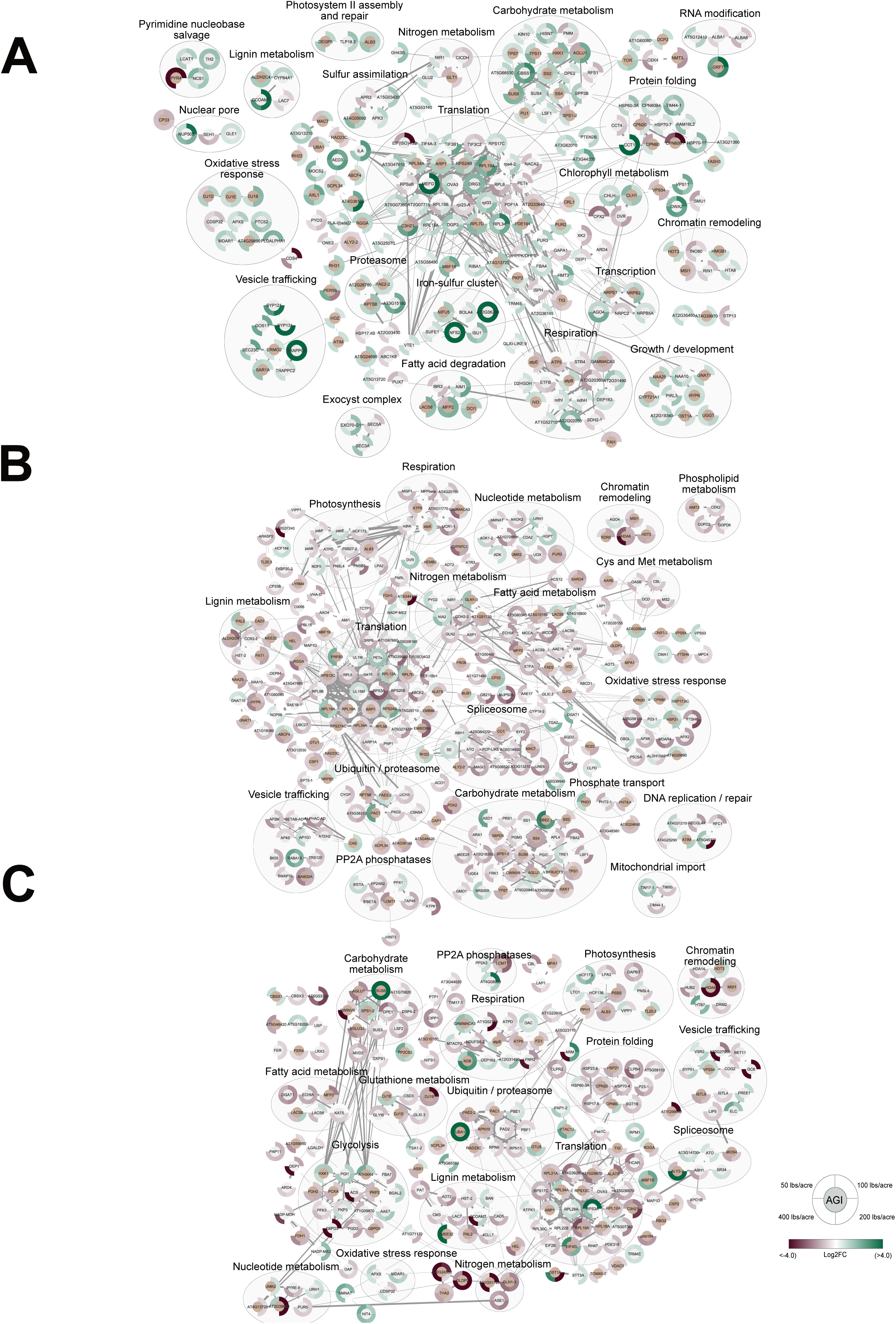
Association network analysis of proteins changing with Humalite addition A-C) STRING-DB association network analysis of significantly changing proteins (q=value <0.05; Fold-change >1.5) with Humalite addition under no urea (A), half urea (B), and full urea (C). Significantly changing proteins were filtered to those appearing in at least 2 Humalite application rates. Wheat gene identifiers were converted to Arabidopsis AGIs for this analysis. Edge thickness indicates strength of connection between nodes. The scale of burgundy to green indicates decrease and increase in abundance, respectively. Groupings of nodes encompassed in a circle indicate proteins involved in the same biological process. Nodes with brown middle indicate proteins common between urea treatments.

## 4. DISCUSSION

An improvement in plant development, growth, and yield with the addition of Humalite has widely been reported for several plant species ^18,26,29,51^. However, with little knowledge about the molecular underpinnings of these responses, especially under field conditions, it remains unclear how Humalite will work for agricultural producers. In parallel with a large-scale field study^19^, we took a quantitative proteomic approach to understand changes in biological processes behind the yield responses observed in field-grown wheat (*T. aestivum*). Our quantitative proteomic results indicate that Humalite induces molecular changes in the abundance of proteins involved in nitrogen assimilation and metabolism, iron-sulfur cluster assembly, carbohydrate and lipid metabolism, oxidative pentose phosphate pathway and glycolysis, along with changes in the abundance of proteins involved in stress response, including those involved in fatty acid beta-oxidation, vesicle trafficking and protein folding / heat shock response.

### 4.1. Humalite induces changes in proteins associated with plant nutrient and energy metabolism

Wheat plants grown with Humalite under the ‘No Urea’ treatment exhibited the maximal molecular benefits, showing an increase in proteins associated with nutrient metabolism, including nitrogen assimilation and nutrient sensing, carbohydrate metabolism and energy metabolism such as enzymes involved in the oxidative pentose phosphate pathway and glycolysis.

#### Nitrogen Metabolism

N is an essential nutrient for plant growth and development, acting as a component of amino acids, building blocks of proteins, nucleic acids, hormones, chlorophyll, and many important metabolites^54^. Our results indicate significant changes in several proteins involved in N metabolism upon Humalite application. A previous study found an increase in the activity of enzymes associated with nitrogen assimilation upon the addition of humic acids, such as NITRATE REDUCTASE, GLUTAMINE SYNTHETASE and GLUTAMATE SYNTHASE, in a dose-dependent manner^55^. Together, GLUTAMINE SYNTHETASE, GLUTAMATE SYNTHASE and GLUTAMATE DEHYDROGENASE form a key link between N and C metabolism whereby nitrate and ammonia are assimilated into amino acids^56^. Previously, NITRATE REDUCTASE and GLUTAMINE SYNTHETASE were found to be up-regulated in response to the addition of humic acids in maize roots^13^. Similarly, we find NITRATE REDUCTASE (NIA2; TraesCS7A02G078500) up-regulated under both ‘No Urea’ and ‘Half Urea’ treatments, showing a positive correlation under ‘Half Urea’ with the amount of Humalite applied.

Further, GLUTAMINE SYNTHETASE, (GLN1;3; TraesCS4B02G240900) was also up-regulated with Humalite addition under ‘No Urea’ and ‘Half Urea’, but inversely correlated with Humalite level under ‘Full Urea’. GLN2 (TraesCS2A02G500400) and the NODULIN GLUTAMINE SYNTHASE (NodGS; TraesCS1A02G143000) were also up-regulated with 448 kg ha^-1^ Humalite under ‘No Urea’ but down-regulated with Humalite addition under ‘Half Urea’. The NADH-dependent glutamate synthase, GLT1 (TRAESCS3A02G266300), and the ferredoxin-dependent glutamate synthase, GLU2 (TRAESCS2B02G152900) were both down-regulated with Humalite under ‘No Urea’, but GLT1 was up-regulated with 224 kg ha^-1^ Humalite under ‘Half Urea’. Finally, GLUTAMATE DEHYDROGENASE2 (GDH2; TraesCS2A02G389900) was up-regulated under ‘No Urea’ with the addition of 448 kg ha^-1^ Humalite but down-regulated with the addition of Humalite under ‘Half Urea’. Together, this suggests that the addition of Humalite induces changes in N assimilation, with a preference for increased N assimilation into glutamine under low N conditions (e.g. ‘No Urea’), but to glutamate, or further amino acid products, with increased N supply (e.g. ‘Half Urea’ or ‘Full Urea’).

#### Carbohydrate Metabolism

Critically important to overall plant metabolism, in particular N assimilation, is the availability of carbohydrates. Several studies have reported changes in starch and sucrose content upon treatment with humic acid^57–59^. However, different crops have shown different responses, with maize leaves showing decreased starch and increased sugar content^58^, and wheat leaf showing increased starch^59^. This indicates potential differences amongst different crops for humic acid-induced changes in carbohydrate metabolism; however, with the source and application conditions of these experiments differing substantially, it can be difficult to draw consensus conclusions. In our field experiment, we identify perturbations in the abundance of proteins associated with starch and sucrose metabolism. This includes starch degradation enzymes: DISPROPORTIONATING ENZYME1 (DPE1; TraesCS2A02G159300), LIMIT DEXTRINASE (LDA; TraesCS7D02G133100), ISOAMYLASE3 (ISA3; TraesCS5A02G248700), STARCH EXCESS4 (SEX4; TraesCS5A02G529100; TraesCS4D02G353800) and LIKE SEX4 2 (LSF2; TraesCS5A02G148200). Similarly, we observe changes in the abundance of proteins associated with starch synthesis. These include STARCH SYNTHASE (SS)1 (TraesCS7A02G120300), SS2 (TraesCS6B02G336400; TraesCS1B02G155700; TraesCS1A02G137200; TraesCS6A02G307800), SS4 (TraesCS1A02G353300), ADP-GLUCOSE PYROPHOSPHORYLASE (APL)1 (TraesCS5B02G484700), APL4 (TraesCS1A02G419600), GRANULE-BOUND STARCH SYNTHASE1 (GBSS1; TraesCS2B02G390700) and PHOSPHOGLUCOSE ISOMERASE1 (PGI1; TraesCS5D02G253700). Collectively, we find both starch synthesis and degradation to be up-regulated with the addition of Humalite under ‘No Urea’, but down-regulated under ‘Half Urea’ and ‘Full Urea’. A similar trend was seen for enzymes involved in sucrose metabolism, including SUCROSE PHOSPHATE SYNTHASE (SPSA1; TraesCS6A02G144800; TraesCS7B02G434300), SUCROSE SYNTHASE (SUS)3 (TraesCS4A02G140000), SUS6 (TraesCS7A02G557600; TraesCS2A02G406700) and TREHALOSE-6-PHOSPHATE SYNTHASE1 (TPS1; TraesCS1A02G339300). Carbohydrate metabolism provides C for use in N metabolism, representing a close connection between the two, and indicating a general trend of Humalite stimulating plant primary metabolism. Although there was not a significant yield increase observed in the field study at this site^19^, this change in enzymes of primary metabolism clearly indicates more effective plant growth. Further, with more urea, the abundance of these enzymes was not stimulated by Humalite, indicating its beneficial impacts are primarily under low N conditions.

#### Nutrient Sensing and Integration

TARGET OF RAPAMYCIN (TOR) kinase and SNF-RELATED KINASE1 (SnRK1) form a signaling module that integrates carbon and nitrogen metabolism^60^. TOR and its regulatory subunits, LST8 and RAPTOR1B, are active under nutrient-rich conditions to promote growth and development through the regulation of ribosome biogenesis and protein synthesis^61,62^.

Antagonistically, the SnRK1 complex is activated under conditions of stress, inducing catabolic processes, such as autophagy, and down-regulating protein synthesis^63^. In line with other studies that observed up-regulation of TOR gene expression upon humic acid treatment^11,44^, we observe increased protein abundance of TOR (TraesCS1A02G227600) with Humalite addition under our ‘No Urea’ treatment. However, with the addition of urea, we do not observe significant changes in TOR protein abundance, except for a down-regulation at the highest Humalite rate. We additionally observe up-regulation of the SnRK1 catalytic subunit, KIN10 (TraesCS3A02G282800), with Humalite application under the ‘No Urea’ treatment, also consistent with several other studies^64,65^. *TOR* gene expression is largely induced by a high cellular nutritional status^66^, such as high amino acid content or nutrient availability through glucose signaling^67^. Interactome and phosphoproteome studies have revealed close ties between SnRK1 and Class II trehalose 6-phosphate synthases (TPS) in both carbon metabolism^68^ and nitrogen metabolism^60^. In Cluster 1 containing TOR and SnRK1, we also identified TPS1 (TraesCS1A02G339300) and the Class II TPS proteins, TPS7 (TraesCS3B02G323900) and TPS11 (TraesCS6A02G351300), as up-regulated with Humalite under ‘No Urea’ and down-regulated under ‘Half Urea’ and ‘Full Urea’ treatments. Humic acids have been shown to interfere with nutrient sensing in *Z. mays* independent of amino acid, sugar and / or organic acid concentrations in both leaf and root^11^. Thus, it is possible that the addition of Humalite under nutrient-poor conditions is interfering with the plant ability to properly sense nutrient availability, disrupting the balance between plant growth and response to conditions of stress.

#### Oxidative Pentose Phosphate Pathway and Glycolysis

Glycolysis and the oxidative pentose phosphate pathway are essential for plant energy metabolism. Within Cluster 1, we observe enrichment of oxidative pentose phosphate pathway enzymes, such as GLUCOSE-6-PHOSPHATE DEHYDROGENASE (G6PD)3 (TraesCS2A02G067800) and G6PD5 (TraesCS2A02G320400), 6-PHOSPHOGLUCOLACTONASE2 (PGL2; TraesCS7A02G270200) and 6-PHOSPHOGLUCONATE DEHYDROGENASE2 (PGD2; TraesCS3B02G565200). Here, we additionally reveal abundance changes in many glycolytic enzymes including: HEXOKINASE1 (HXK1; TraesCS1D02G344600), the cytosolic GLUCOSE-6-PHOSPHATE ISOMERASE (PGIC; TraesCS1A02G040600), FRUCTOSE-BISPHOSPHATE ALDOLASE (FBA)2 (TraesCS4B02G109900), FBA4 (TraesCS3D02G384000), FBA7 (TraesCS7B02G283000), PHOSPHOGLUCOMUTASE3 (PGM3; TraesCS4D02G047300), and PLASTIDIAL PYRUVATE KINASE2 (PKP2; TraesCS3A02G239300). This supports previous research in maize roots treated with humic acids, where an up-regulation of the activity of GLUCOKINASE, PHOSPHOGLUCOSE ISOMERASE, PHOSPHOFRUCTOKINASE, and PYRUVATE KINASE has been reported^69^. Furthermore, an increased abundance of proteins involved in glycolysis and the oxidative pentose phosphate pathway has also been identified in Arabidopsis roots upon humic acid application^47^. Thus, in soils without added urea, Humalite application appears to stimulate energy metabolism, however, upon the addition of urea, Humalite suppresses the abundance of glycolytic and oxidative pentose phosphate pathway enzymes (**Figure 4**). Thus, Humalite appears to affect the overall plant energy metabolism most efficiently under low nutrient conditions.

**Figure 4.**
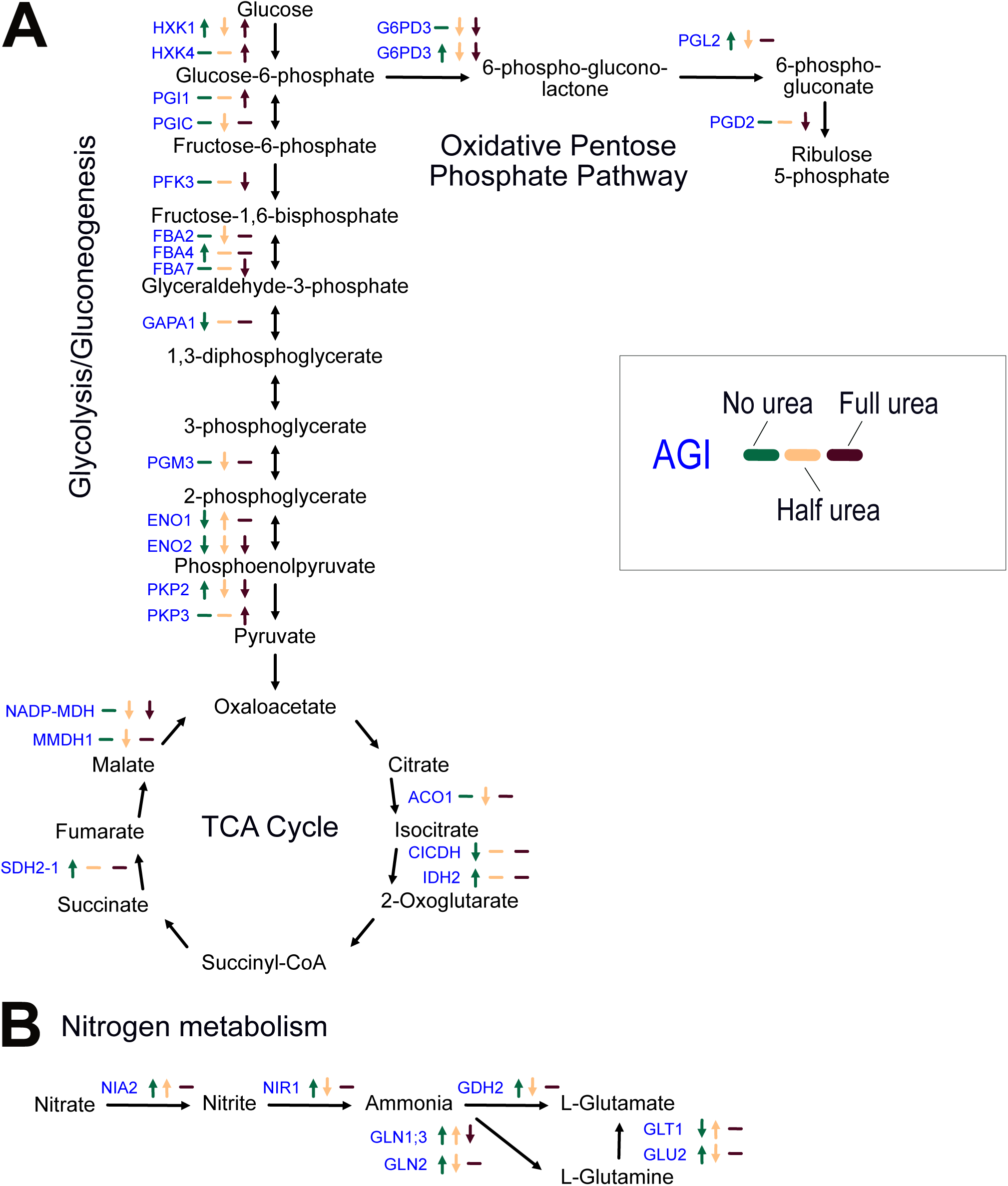
Metabolic pathway changes with addition of Humalite A) Changes in glycolysis/gluconeogenesis, oxidative pentose phosphate pathway and TCA cycle protein abundance with Humalite. B) Changes in protein abundance associated with nitrogen metabolism and assimilation. Arrows indicate direction of protein abundance whether up-regulated or down-regulated. Line indicates no abundance change. Green indicates the change under ‘No Urea’, beige under ‘Half Urea’ and burgundy under ‘Full Urea’. Blue indicates the common name of proteins changing in abundance within our dataset.

#### Iron-Sulfur Cluster Assembly

Closely linked with the increased abundance of energy and nutrient metabolic enzymes with Humalite with the ‘No Urea’ application treatment, Cluster 1 was highly enriched for proteins involved in iron-sulfur (Fe-S) cluster assembly. Fe-S clusters are cofactors for several key plant processes including electron transfer, redox reactions, enzyme catalysis, and sensing iron and oxygen levels^70^. Humic acids have been suggested to form stable complexes with Fe(III), which could improve its biological availability to plants^71^. In our study, we observe increased abundance of the Fe-S cluster assembly proteins LYR MOTIF CONTAINING4 (LYRM4; TraesCS4A02G084700), IRON-SULFUR CLUSTER ASSEMBLY PROTEIN1 (ISU1; TraesCS1A02G406300), NiFU DOMAIN PROTEIN5 (NIFU5; TraesCS1A02G097200), BolA-LIKE4 (BOLA4; TraesCS7A02G232700), SULFURE1 (SUFE1; TraesCS5D02G176300), and NIFS-LIKE CYSTEINE DESULFURASE2 (NFS2; TraesCS4D02G120100) with Humalite addition under the ‘No Urea’ treatment. Further, we also observe the iron transporter OLIGOPEPTIDE TRANSPORTER3 (OPT3; TraesCS5A02G391000) within Cluster 1, showing up-regulation with Humalite under the ‘No Urea’ treatment. Taken together, this indicates that Humalite may promote iron uptake and assimilation under low nutrient conditions. Additionally, humic acids have been reported to enhance electron transport in photosystem II, thereby increasing photosynthetic efficiency^72^, which may be a result of the increased assembly of these iron-sulfur clusters. Thus, Humalite appears to promote plant energy metabolism and usage of nutrients such as C, N and S under low nutrient conditions.

### 4.2. Humalite induces changes in proteins involved with stress response

Humic acids have previously been reported as a promoter of the weak acid stress response when applied to maize seedlings, acting as a priming agent for enhanced plant response to abiotic stressors^64,73^. The addition of humic acids has also been described as inducing a eustress state in rice plants, where an initial stress state leads to enhanced photosynthetic and metabolic performance as a sort of priming mechanism^74^. In addition to the altered metabolic landscape, upon the addition of Humalite in wheat across a range of urea treatments, we observed the induction of several proteome changes related to plant stress response(s), including changes in vesicle trafficking and chaperone proteins.

#### Vesicle Trafficking

Proteins involved in vesicle and protein transport were also highly enriched in Cluster 1. The COP-II coat is involved in vesicle secretion from the ER and its coat-GTPase, SAR1 recruits COP-II coat proteins^75^. In Cluster 1 of our dataset, we observe the COP-II coat proteins SEC23C (TraesCS5D02G247600) and SEC24A (TraesCS2B02G008900) and the SAR1 ARF-like COP-II coat GTPase, SAR1C (TraesCS3A02G172900) as well as the coat-like ESCRT-III protein, VPS2 (TraesCS4A02G174700). The Rab family of small GTPases are involved in motor protein recruitment, cargo selection and docking factor interaction in eukaryotes. Cluster 1 included the wheat RABA2C (TraesCS5A02G485100) and the PRENYLATED RAB ACCEPTOR (PRA1B2; TraesCS3A02G348100).

Additionally, we observe the Arl8-like small GTPase, ARL8A (TraesCS6A02G293900). Finally, we observe several SNARE proteins within Cluster 1, including the Qa SNAREs: SYP121 (TraesCS5A02G425600), SYP131 (TraesCS2A02G246600), the Qb SNARE GOS11 (TraesCS5A02G226000), the Qc SNARE SYP51 (TraesCS5A02G062300), and the R-SNARE VAMP712 (TraesCS7A02G136300). Although vesicle trafficking has been shown in other proteomic studies to be influenced by humic acid addition^13, 47^, we did not observe the same vesicle trafficking-associated proteins as were previously identified. Thus, our study may have preferentially identified proteins up-regulated by Humalite under low nutrient conditions.

#### Protein Folding

Several heat shock proteins and chaperone proteins were up-regulated within our dataset with Humalite application, validating previous studies of the effects of Humalite on roots^13,47^. Interestingly, humic acids have also been surmised to improve abiotic stress tolerance, such as heat shock, through the up-regulation of several heat shock protein genes^35^, while we observe this at the protein-level. Specifically, we found heat shock proteins to be up-regulated with Humalite under ‘No Urea’ and ‘Half Urea’ treatments, but down-regulated under ‘Full Urea’. Similar to their abiotic stress responses, our results may also be indicative of a stress response priming mechanism, further suggesting that humic acids and Humalite act as a eustress in wheat^74^.

### 4.3. Conclusions

We present a comprehensive analysis of the proteome changes elicited in field-grown wheat grown under different Humalite application rates with urea fertilizer applied at the fully recommended or half recommended rate based on soil testing prior to sowing, including a no Humalite and no urea control. We find that most biological processes are up-regulated with Humalite under a ‘No Urea’ growth treatment, whereas many of the same processes are down-regulated upon urea application. These results collectively suggest that Humalite is more effective under N limiting growth conditions, which is an ideal scenario for improving crop N-use efficiency. This hypothesis is supported by extensive up-regulation of proteins involved in nitrogen, carbon, and sulfur metabolism as well as photosynthesis and the electron transport chain. In addition, we observe potential priming through up-regulation of plant stress responsive processes, such as vesicle trafficking and heat shock proteins. Overall, we present a comprehensive proteomic analysis of the molecular mechanisms governing growth promotion by Humalite, with and without urea. This resource can inform agricultural producers on the optimal application regimes for usage of Humalite and urea as a growth stimulant.

## Supporting information

Supplemental Tabl 2

## ACKNOWLEDGEMENTS

The authors thank MITACs and Canada Foundation for Innovation (CFI) for funding. The authors also thank Jack Moore of the Alberta Proteomics and Mass Spectrometry Facility for assistance with mass spectrometer operation and maintenance. Finally, the authors thank WestMet Ag for co-funding the MITACs Accelerate project and for providing Humalite.

## AUTHOR CONTRIBUTIONS

LEG: Investigation; Formal Analysis; Methodology; Data curation & Visualisation, Writing & Editing.

MT: Investigation; Methodology.

LYG: Conceptualization; Field Trial; Wheat Growth and Harvesting; Reviewing & Editing; Funding Acquisition.

RGU: Conceptualization; Methodology; Supervision; Project Administration; Data curation; Writing & Editing; Funding Acquisition.

## DATA AVAILABILITY

The datasets that we have presented in this study are available in public repositories. All proteomic data has been submitted to the PRoteomics IDEntifications Database (PRIDE; https://www.ebi.ac.uk/pride/) with the identification number PXD058950.

---

**Supp Figure 1.**
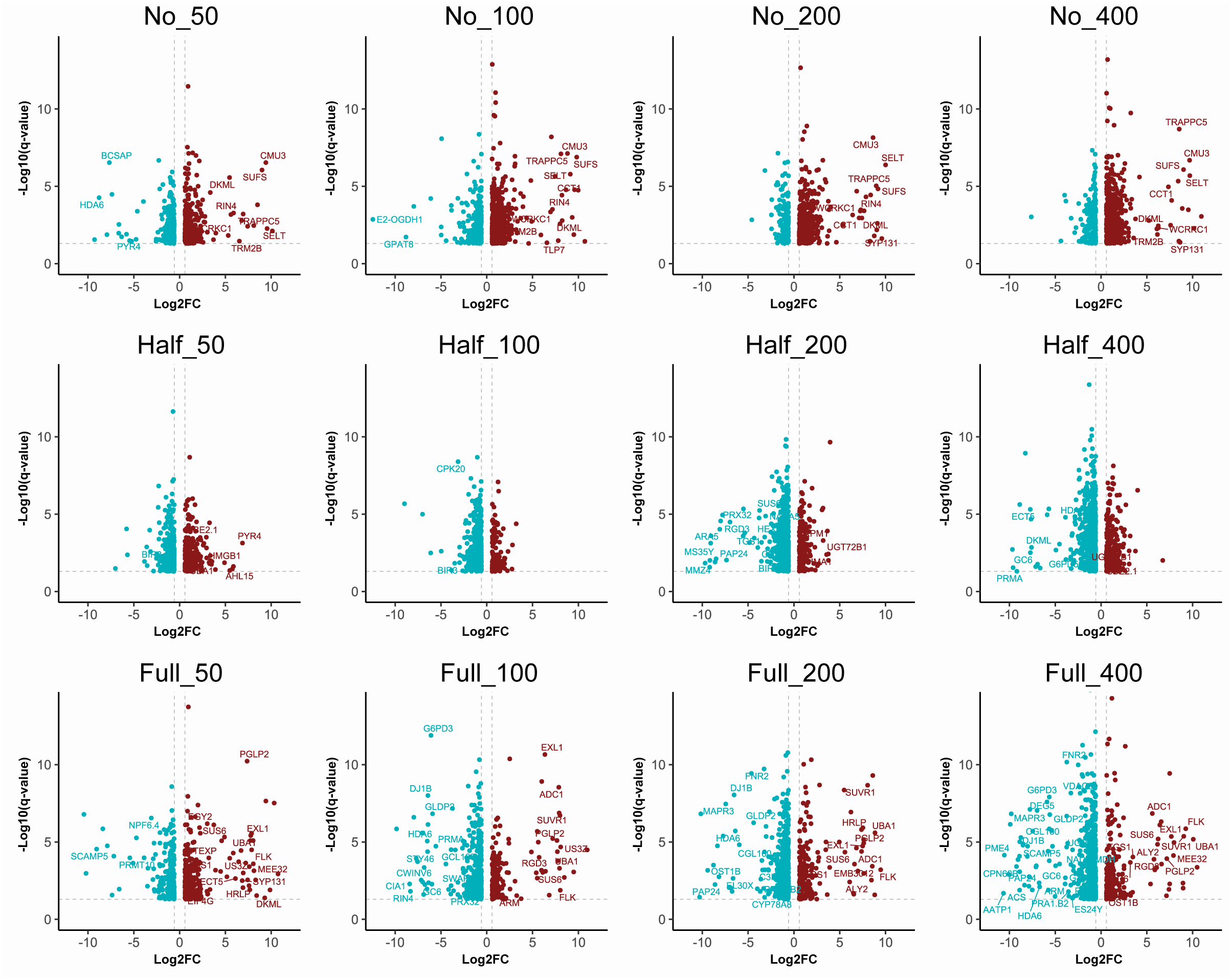
Volcano plots of Humalite under different urea concentrations Volcano plots of significantly changing proteins (q-value < 0.05; Fold change > 1.5) under No Urea, Half Urea and Full Urea conditions with added Humalite (50, 100, 200, 400 kg/ha). Burgundy represents up-regulated proteins and blue represents down-regulated proteins.

## SUPPLEMENTARY DATA

**Supp Table 1.** Summary of all quantified and significantly changing proteins

**Supp Table 2.** AraCyc metabolic pathway analysis

